## Supplementary material for "Distinct trafficking routes of polarized and non-polarized membrane cargoes in *Aspergillus nidulans*"

Key words: fungi; traffic; sorting; COPII; Golgi; secretion; SNAREs

Running title: Protein trafficking via Golgi-bypass

### Supplementary Material

**Supplementary Figure S1**. 7 min video of *de novo* UapA-GFP *versus* mCherry-SynA after 3h of derepression. Maximal intensity projection of deconvolved video showing that cytoplasmic UapA and SynA structures do not colocalize significantly (PCC=0.01, n=52) apart from a small number of doubly labeled puncta (0.33±0.03 μm) with low-range mobility. Scale bar: 5μm

Link: Figure S1.mp4

**Supplementary Figure S2**. 3 min video of *de novo* UapA-GFP trafficking dynamics after 3h of derepression. Maximal intensity projection of deconvolved video showing cytoplasmic oscillating thread structures decorated by pearl-like foci (0.32±0.02 μm) and a faint vesicular/tubular network. Scale bar: 5μm

Link: Figure S2.mp4

**Supplementary Figure S3**. 2 min video of *de novo* mCherry-SynA after 3h of derepression. Maximal intensity projection of deconvolved video showing thread like structures with pearls and puncta moved towards the apical area. Scale bar: 5μm

Link: Figure S3.mp4

**Supplementary Figure S4**. 10 min video of mCherry-SedV *versus de novo* UapA-GFP after 3h of derepression. Maximal intensity projection of deconvolved video showing that neosynthesized UapA does not colocalize significantly with the ERGIC/early-Golgi marker SedV (PCC=0.01, n=100). Scale bar: 5μm

Link: Figure S4.mp4

**Supplementary Figure S5**. 6 min video of mRFP-PH$^{OSBP}$ *versus de novo* UapA-GFP after 3h of derepression. Maximal intensity projection of deconvolved video showing that neosynthesized UapA does not colocalize with the late-Golgi marker PH$^{OSBP}$ (PCC=0.01, n=35). Scale bar: 5μm

Link: Figure S5.mp4

**Supplementary Figure S6**. 5 min video of mCherry-SedV *versus de novo* GFP-SynA after 3h of derepression. Maximal intensity projection of deconvolved video showing that neosynthesized SynA does not colocalize significantly with the ERGIC/early-Golgi marker SedV (PCC=0.04, n=40). Scale bar: 5µm

Link: Figure S6.mp4

**Supplementary Figure S7**. 6 min video of mRFP-$PH^{osbp}$ *versus de novo* GFP-SynA after 3h of derepression. Maximal intensity projection of deconvolved video showing that neosynthesized SynA colocalizes significantly with the late-Golgi marker $PH^{osbp}$ (PCC=0.74, n=76, p<0.0001). Scale bar: 5µm

Link: Figure S7.mp4

**Supplementary Figure S8**. Super Resolution Radial Fluctuation (SRRF) imaging of colocalization of SynA with the late Golgi marker $PH^{osbp}$ in fixed cells. Maximal intensity projection of 57 z-stacks (left panel) and a specific z-stack (z = 8, right panel) showing a significant colocalization of GFP-SynA with the late-Golgi marker mRFP-$PH^{OSBP}$.

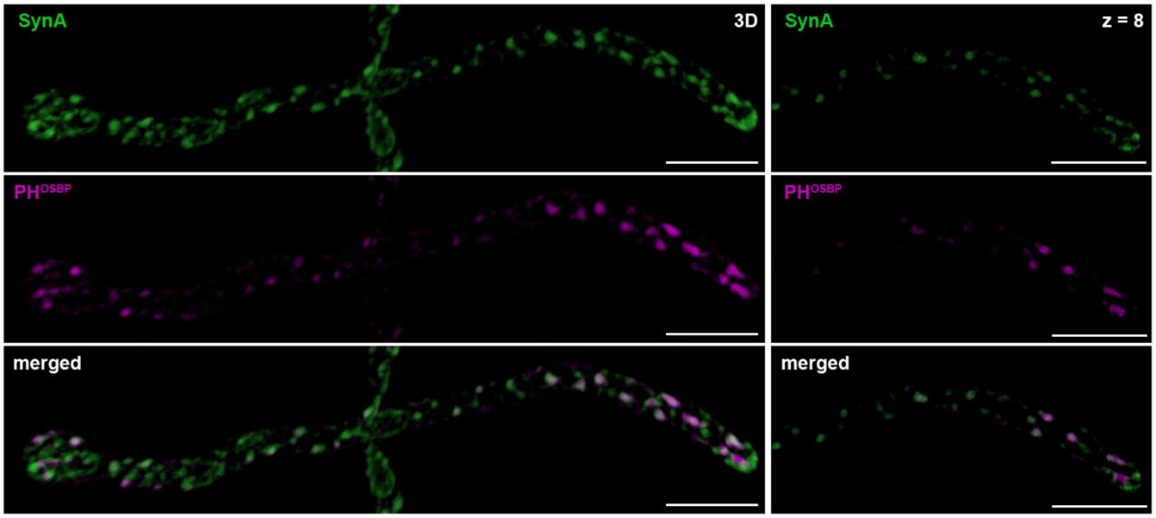

**Supplementary Figure S9**. UapA membrane aggregates upon RabE repression do not colocalize with early- or late-Golgi markers. Maximal intensity projections of deconvolved z-stacks, using *thiA<sub>p</sub>-rabE* strains co-expressing UapA-GFP and the early Golgi marker mCherry-SedV (left panel) or the late Golgi marker mRFP-$PH^{osbp}$ (right panel). Notice the almost non-existent degree of overlap (white color in the merged image) between UapA-GFP and Golgi-markers. Scale bars: 5μm. Quantification of co-localization by calculating Pearson's correlation coefficient (PCC) verifies the low degree of co-localization of UapA with both early- and late-Golgi markers (PCC=0.17±0.07, n=6 with SedV and PCC=0.26±0.08, n=11 with $PH^{osbp}$).

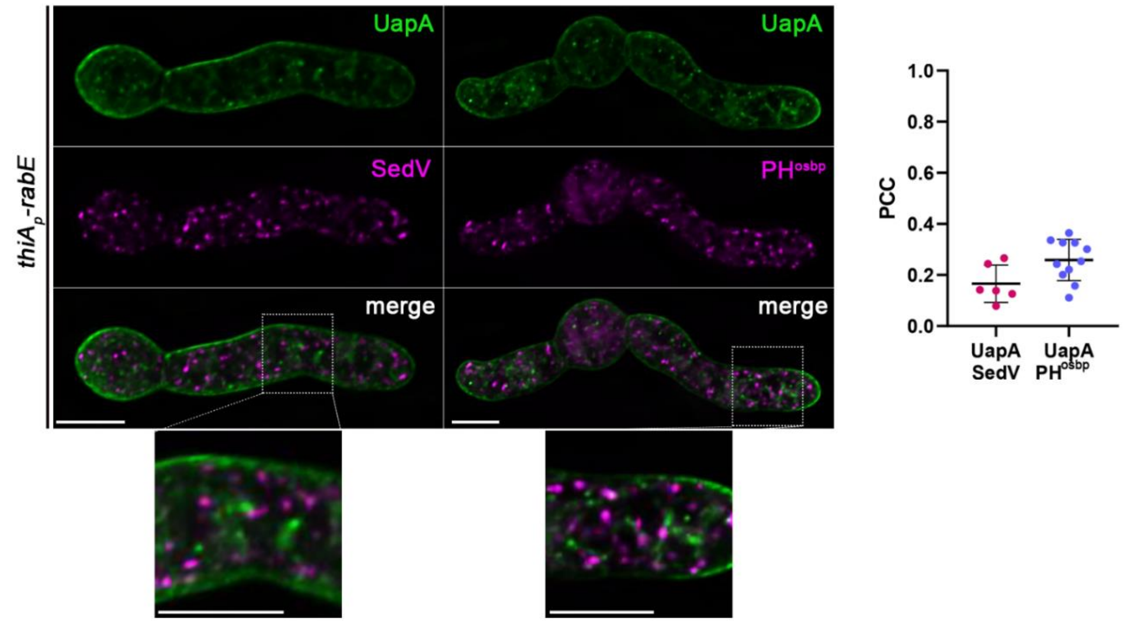


**Supplementary Figure S10**. Triple R-SNARE knockout Δ*sec22* Δ*nyvA* Δ*synA* does not affect UapA or ChsB trafficking to the PM. (A) Growth phenotypes of null mutants of R-SNAREs at 37°C and 42°C, compared to an isogenic wild-type (wt) control strain. Δ*nyvA* shows no growth defect, while the triple knock-out Δ*sec22* Δ*nyvA* Δ*synA* exhibits a severe growth defect at both temperatures. (B) Maximal intensity projections of deconvolved z-stacks, using strains expressing UapA-GFP or GFP-ChsB in genetic backgrounds where single or triple R-SNAREs are deleted. Deletion of either *nyvA* or all three R-SNAREs has no effect on the trafficking of both cargoes to the PM. Scale bars:

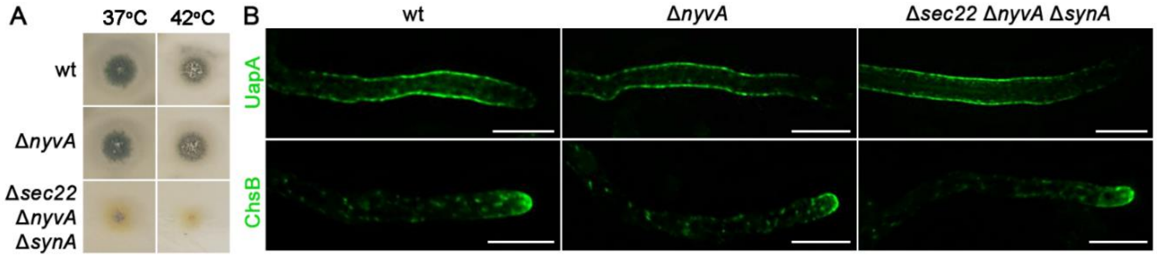

5µm

**Supplementary Table 1. Strains used in this study**

All strains carry the *veA1* mutation affecting sporulation. *pabaA1, pyroA4, riboB2, argB2, pyrG89, and inoB2* are auxotrophic mutations for p-aminobenzoic acid, pyridoxine, riboflavin, arginine, uracil/uridine and inositol respectively. *wA4* is a mutation resulting in white conidiospore color.

| **Name** | **Genotype** | **Reference** |
|---|---|---|
| TNO2A7 | *ΔnkuA::argB pyrG89 pyroA4 riboB2* | Nayak et al., 2006 |
| wt | *pabaA1* | wild-type reference strain |
| *$alcA_p$-uapA-gfp $alcA_p$-mCherry-synA* | *ΔuapA::$alcA_p$-uapA-GFP::AFriboB $alcA_p$-mCherry-synA::AFpyroA ΔnkuA::argB riboB2 pyroA4 pabaA1* | This study |
| Δ*uapA* | *ΔuapA riboB2* | Pantazopoulou et al., 2009 |
| Δ*synA* | *ΔsynAΔ::AFriboB ΔnkuA::argB pyrG89 riboB2 pyroA4* | This study |
| *$alcA_p$-uapA-gfp mCherry-sedV* | *pyroA::$gpdA^m_p$::mCherry-sedV ΔuapA::$alcA_p$-uapA-gfp::AFriboB pabaA1 inoB2* | Dimou et al., 2020 |
| *$alcA_p$-uapA-gfp mRFP-$PH^{OSBP}$* | *pyroA::$gpdA^m_p$::mRFP-$PH^{OSBP}$ ΔuapA::$alcA_p$-uapA-gfp::AFriboB pabaA1 inoB2* | Dimou et al., 2020 |
| *$alcA_p$-gfp-synA mCherry-sedV* | *pyroA::$gpdA^m_p$::mCherry-sedV $alcA_p$-gfp-synA::AFpyrG wA4 inoB2* | Dimou et al., 2020 |
| *$alcA_p$-gfp-synA mrfp-$PH^{OSBP}$* | *pyroA::$gpdA^m_p$::mrfp-$PH^{OSBP}$ $alcA_p$-gfp-synA::AFpyrG wA4 inoB2* | Dimou et al., 2020 |
| *$thiA_p$-sarA $alcA_p$-uapA-gfp $alcA_p$-mCherry-synA* | *ΔuapA::$alcA_p$-uapA-gfp::AFriboB $alcA_p$-mCherry-synA::AFpyroA $thiA_p$-sarA::AFpyrG ΔnkuA::argB riboB2 pyroA4* | This study |
| *$thiA_p$-sec12 $alcA_p$-uapA-gfp $alcA_p$-mCherry-synA* | *ΔuapA::$alcA_p$-uapA-gfp::AFriboB $alcA_p$-mCherry-synA::AFpyroA $thiA_p$-sec12::AFriboB ΔnkuA::argB riboB2 pyroA4 pabaA1* | This study |
| *$thiA_p$-sec24 $alcA_p$-uapA-gfp $alcA_p$-mCherry-synA* | *ΔuapA::$alcA_p$-uapA-GFP::AFriboB $alcA_p$-mCherry-synA::AFpyroA $thiA_p$-sec24::AFpyrG ΔnkuA::argB riboB2 pyroA4* | This study |
| *$thiA_p$-sec13 $alcA_p$-uapA-gfp $alcA_p$-mCherry-synA* | *ΔuapA::$alcA_p$-uapA-GFP::AFriboB $alcA_p$-mCherry-synA::AFpyroA $thiA_p$-sec13::AFpyrG ΔnkuA::argB riboB2 pyroA4* | This study |
| *$thiA_p$-sec31 $alcA_p$-uapA-gfp $alcA_p$-mCherry-synA* | *ΔuapA::$alcA_p$-uapA-GFP::AFriboB $alcA_p$-mCherry-synA::AFpyroA $thiA_p$-sec31::pabaA ΔnkuA::argB riboB2 pyroA4* | This study |
| *$thiA_p^{FLAG}$-sarA* | *$thiA_p^{FLAG}$-sarA::AFpyrG ΔnkuA::argB pyrG89 pyroA4 riboB2* | This study |
| *$thiA_p^{FLAG}$-sec12* | *$thiA_p^{FLAG}$-sec12::AFpyrG ΔnkuA::argB pyrG89 pyroA4 riboB2* | This study |
| *$thiA_p^{FLAG}$-sec24* | *$thiA_p^{FLAG}$-sec24::AFpyrG ΔnkuA::argB pyrG89 pyroA4 riboB2* | This study |

| Name | Genotype | Reference |
|---|---|---|
| *thiA$_p$$^{FLAG}$-sec13* | *thiA$_p$$^{FLAG}$-sec13::AFpyrG ΔnkuA::argB pyrG89 pyroA4 riboB2* | This study |
| *thiA$_p$$^{FLAG}$-sec31* | *thiA$_p$$^{FLAG}$-sec31::AFpyrG ΔnkuA::argB pyrG89 pyroA4 riboB2* | This study |
| *sec31$^{ts}$-AV* | *sec31A1249V::AFpyrG ΔnkuA::argB pyrG89 riboB2 pyroA4* | This study |
| *sec31$^{ts}$-AS* | *sec31A1249S::AFpyrG ΔnkuA::argB pyrG89 riboB2 pyroA4* | This study |
| *sec31$^{ts}$-AG* | *sec31A1249G::AFpyrG ΔnkuA::argB pyrG89 riboB2 pyroA4* | This study |
| *sec31$^{ts}$-AP* | *sec31A1249P::AFpyrG ΔnkuA::argB pyrG89 riboB2 pyroA4* | This study |
| *sec31$^{ts}$-AP alcA$_p$-uapA-gfp alcA$_p$-mCherry-synA* | *sec31A1249P::AFpyrG ΔuapA::alcA$_p$-uapA-gfp::AFriboB alcA$_p$-mCherry-synA::AFpyroA ΔnkuA::argB riboB2 pyroA4* | This study |
| *sec31$^{ts}$-AP alcA$_p$-uapA-gfp sec16-mCherry* | *sec31A1249P::AFpyrG ΔuapA::alcA$_p$-uapA-gfp::AFriboB sec16-mCheery::AFpyroA ΔnkuA::argB riboB2 pyroA4* | This study |
| *sec31$^{ts}$-AP alcA$_p$-gfp-synA sec16-mCherry* | *sec31A1249P::AFpyrG ΔuapA::alcA$_p$-gfp-synA::AFriboB sec16-mCheery::AFpyroA ΔnkuA::argB riboB2 pyroA4* | This study |
| *thiA$_p$-copA alcA$_p$-uapA-gfp alcA$_p$-mCherry-synA* | *ΔuapA::alcA$_p$-uapA-gfp::AFriboB alcA$_p$-mCherry-synA::AFpyroA thiA$_p$-copA::AFpyrG ΔnkuA::argB riboB2 pyroA4* | This study |
| *thiA$_p$-arfA alcA$_p$-uapA-gfp alcA$_p$-mCherry-synA* | *ΔuapA::alcA$_p$-uapA-gfp::AFriboB alcA$_p$-mCherry-synA::AFpyroA thiA$_p$-arfA::AFpyrG ΔnkuA::argB riboB2 pyroA4* | This study |
| *thiA$_p$-sedV alcA$_p$-uapA-gfp alcA$_p$-mCherry-synA* | *ΔuapA::alcA$_p$-uapA-gfp::AFriboB alcA$_p$-mCherry-synA::AFpyroA thiA$_p$-sedV::AFpyrG ΔnkuA::argB riboB2 pyroA4 pabaA1* | This study |
| *thiA$_p$-geaA alcA$_p$-uapA-gfp alcA$_p$-mCherry-synA* | *ΔuapA::alcA$_p$-uapA-gfp::AFriboB alcA$_p$-mCherry-synA::AFpyroA thiA$_p$-geaA::AFpyrG ΔnkuA::argB riboB2 pyroA4 pabaA1* | This study |
| *thiA$_p$-rabO alcA$_p$-uapA-gfp alcA$_p$-mCherry-synA* | *ΔuapA::alcA$_p$-uapA-gfp::AFriboB alcA$_p$-mCherry-synA::AFpyroA thiA$_p$-rabO::AFpyrG ΔnkuA::argB riboB2 pyroA4 pabaA1* | This study |
| *thiA$_p$-hypB alcA$_p$-uapA-gfp alcA$_p$-mCherry-synA* | *ΔuapA::alcA$_p$-uapA-gfp::AFriboB alcA$_p$-mCherry-synA::AFpyroA thiA$_p$-hypB::AFpyrG ΔnkuA::argB riboB2 pyroA4* | This study |
| *thiA$_p$-rabE alcA$_p$-uapA-gfp alcA$_p$-mCherry-synA* | *ΔuapA::alcA$_p$-uapA-gfp::AFriboB alcA$_p$-mCherry-synA::AFpyroA thiA$_p$-rabE::AFpyrG ΔnkuA::argB riboB2 pyroA4* | This study |
| *thiA$_p$-ap1$^σ$ alcA$_p$-uapA-gfp alcA$_p$-mCherry-synA* | *ΔuapA::alcA$_p$-uapA-gfp::AFriboB alcA$_p$-mCherry-synA::AFpyroA thiA$_p$-ap1$^σ$::AFriboB ΔnkuA::argB riboB2 pyroA4 pabaA1* | This study |
| *alcA$_p$-uapA-gfp mCherry-sedV thiA$_p$-rabE* | *ΔuapA::alcA$_p$-uapA-gfp::AFriboB pyroA::gpdA$^m$$_p$::mCherry-sedV thiA$_p$-rabE::AFpyrG* | This study |
| *thiA$_p$-rabE alcA$_p$-uapA-gfp* | *pyroA::gpdA$^m$$_p$::mRFP-PH$^{OSBP}$ ΔuapA::alcA$_p$-* | This study |

| Name | Genotype | Reference |
|---|---|---|
| *mRFP-PH$^{OSBP}$* | *uapA-gfp::AFriboB thiA$_p$-rabE::AFpyrG pabaA1* | |
| *thiA$_p$-ykt6 alcA$_p$-uapA-gfp alcA$_p$-mCherry-synA* | *ΔuapA::alcA$_p$-uapA-gfp::AFriboB alcA$_p$-mCherry-synA::AFpyroA thiA$_p$-ykt6::AFpyrG ΔnkuA::argB riboB2 pyroA4 pabaA1* | This study |
| *Δsec22 alcA$_p$-uapA-gfp alcA$_p$-mCherry-synA* | *ΔuapA::alcA$_p$-uapA-gfp::AFriboB alcA$_p$-mCherry-synA::AFpyroA Δsec22::AFpyrG ΔnkuA::argB riboB2 pyroA4 pabaA1* | This study |
| *thiA$_p$-ykt6 Δsec22 alcA$_p$-uapA-gfp alcA$_p$-mCherry-synA* | *ΔuapA::alcA$_p$-uapA-gfp::AFriboB alcA$_p$-mCherry-synA::AFpyroA thiA$_p$-ykt6::AFriboB Δsec22::AFpyrG ΔnkuA::argB riboB2 pyroA4* | This study |
| *thiA$_p$-sft1 alcA$_p$-uapA-gfp alcA$_p$-mCherry-synA* | *ΔuapA::alcA$_p$-uapA-gfp::AFriboB alcA$_p$-mCherry-synA::AFpyroA thiA$_p$-sft1::pabaA ΔnkuA::argB riboB2 pyroA* | This study |
| *thiA$_p$-bos1 alcA$_p$-uapA-gfp alcA$_p$-mCherry-synA* | *ΔuapA::alcA$_p$-uapA-gfp::AFriboB alcA$_p$-mCherry-synA::AFpyroA thiA$_p$-bos1::pabaA ΔnkuA::argB riboB2 pyroA4* | This study |
| *thiA$_p^{FLAG}$-ykt6* | *thiA$_p^{FLAG}$-ykt6::pyrG89 ΔnkuA::argB pyrG89 pyroA4 riboB2* | This study |
| *thiA$_p^{FLAG}$-sft1* | *thiA$_p^{FLAG}$-sft1::AFpyrG ΔnkuA::argB pyrG89 pyroA4 riboB2* | This study |
| *thiA$_p^{FLAG}$-bos1* | *thiA$_p^{FLAG}$-bos1::AFpyrG ΔnkuA::argB pyrG89 pyroA4 riboB2* | This study |
| *ΔrabD* | *ΔrabD::AFpyrG ΔnkuA::argB pyrG89 riboB2 pyroA4* | This study |
| *thiA$_p$-ssoA* | *thiA$_p$-ssoA::AFpyrG ΔnkuA::argB pyrG89 riboB2 pyroA4* | Dimou et al., 2020 |
| *thiA$_p$-sec9* | *thiA$_p$-sec9::AFpyrG ΔnkuA::argB pyrG89 riboB2 pyroA5* | This study |
| *thiA$_p^{FLAG}$-ssoA* | *thiA$_p^{FLAG}$-ssoA::AFpyrG ΔnkuA::argB pyrG89 pyroA4 riboB2* | This study |
| *thiA$_p^{FLAG}$-sec9* | *thiA$_p^{FLAG}$-sec9::AFpyrG ΔnkuA::argB pyrG89 pyroA4 riboB2* | This study |
| *uapA-gfp* | *ΔuapA::uapA-gfp::AFriboB ΔuapC::AfpyrG ΔnkuA::argB pabaA1 pyroA4 riboB2* | Evangelinos et al., 2016 |
| *thiAp-ssoA uapA-gfp* | *ΔuapA::uapA-gfp thiA$_p$-ssoA::AFpyrG ΔnkuA::argB pabaA1* | Dimou et al., 2020 |
| *thiAp-sec9 uapA-gfp* | *ΔuapA::uapA-gfp thiA$_p$-sec9::AFpyrG ΔnkuA::argB pabaA1* | This study |
| *ΔsynA uapA-gfp* | *ΔuapA::uapA-gfp ΔsynA:AFriboB ΔnkuA::argB pabaA1* | This study |
| *gfp-chsB* | *gfp-chsB::AFpyrG ΔnkuA::argB pyrG89 pyroA4 riboB2* | Dimou et al., 2020 |
| *thiAp-ssoA gfp-chsB* | *gfp-chsB::AFpyrG thiA$_p$-ssoA::AFriboB ΔnkuA::argB pyrG89 pyroA4 riboB2* | This study |
| *thiAp-sec9 gfp-chsB* | *gfp-chsB::AFpyrG thiA$_p$-sec9::AFpyrG pyrG89 pyroA4* | This study |
| *ΔsynA gfp-chsB* | *gfp-chsB::AFpyrG ΔsynA::AFriboB ΔnkuA::argB pyrG89 pyroA4 riboB2* | This study |

| Name | Genotype | Reference |
|---|---|---|
| *ΔrabD alcA*$_p$*-uapA-gfp alcA*$_p$*-mCherry-synA* | *ΔuapA::alcA*$_p$*-uapA-gfp::AFriboB alcA*$_p$*-mCherry-synA::AFpyroA ΔrabD::AFpyrG ΔnkuA::argB riboB2 pyroA4 pabaA1* | This study |
| *ΔnyvA* | *ΔnyvA::AFpyrG ΔnkuA::argB pyrG89 riboB2 pyroA4* | This study |
| *Δsec22 ΔnyvA ΔsynA* | *Δsec22::AFpyrG ΔnyvA::AFpyroA ΔsynA::AFriboB ΔnkuA::argB pyrG89 riboB2 pyroA4* | This study |
| *ΔnyvA uapA-gfp* | *ΔuapA::uapA-gfp ΔnyvA:AFpyrG ΔnkuA::argB pabaA1* | This study |
| *Δsec22 ΔnyvA ΔsynA uapA-gfp* | *ΔuapA::uapA-gfp Δsec22::AFpyrG ΔnyvA::AFpyrG ΔsynA::AFriboB ΔnkuA::argB pyroA4 pabaA1* | This study |
| *ΔnyvA gfp-chsB* | *gfp-chsB::AFpyrG ΔnyvA:AFpyrG pyrG89 pyroA4* | This study |
| *Δsec22 ΔnyvA ΔsynA gfp-chsB* | *gfp-chsB::AFpyrG Δsec22::AFpyrG ΔnyvA::AFpyrG ΔsynA::AFriboB ΔnkuA::argB pyroA4 pabaA1* | This study |

**Supplementary Table 2. Annotation of proteins used in this study**

| FungiDB ID | Systematic/Protein name | Trafficking process |
|---|---|---|
| AN0411 | SarA$^{Sar1}$ | ER exit sites (ERes) |
| AN3720 | Sec24 | ER exit sites (ERes) |
| AN4317 | Sec13 | ER exit sites (ERes) |
| AN6257 | Sec31 | ER exit sites (ERes) |
| AN11127 | Sec12 | ER exit sites (ERes) |
| AN3026 | CopA$^{CopI}$ | COPI coat |
| AN1126 | ArfA$^{Arf1/2}$ | Early/late Golgi sorting |
| AN0112 | GeaA$^{Gea1/2}$ | Early Golgi sorting |
| AN6709 | HypB$^{Sec7}$ | Late Golgi sorting |
| AN4281 | RabO$^{Ypt1/Rab1}$ | Early Golgi sorting |
| AN0347 | RabE$^{Ypt31/32/Rab11}$ | Post-Golgi |
| AN7682 | Ap1$_{\sigma}^{Aps1}$ | Late Golgi sorting |
| AN6974 | RabD$^{Sec4}$ | exocyst |
| AN8488 | Ykt6 | R-SNARE |
| AN8769 | SynA$^{Snc1/2}$ | R-SNARE |
| AN0571 | NyvA$^{Nyv1}$ | R-SNARE |
| ChrI_A_nidulans_FGSC_A4 2,596,682-2,597,694 | Sec22 | R-SNARE |
| AN3416 | SsoA$^{Sso1/2}$ | Qa-SNARE |
| AN9526 | SedV$^{Sed5}$ | Qa-SNARE |
| AN2419 | Sec9 | Qb-SNARE |

| FungiDB ID | Systematic/Protein name | Trafficking process |
|---|---|---|
| AN11900 | Bos1 | Qb-SNARE |
| AN10508 | Sft1 | Qc-SNARE |
| AN2523 | ChsB$^{Chs3}$ | apical cargo |
| AN6932 | UapA | non-polar cargo |

**Supplementary Table 3. Primers used for cloning and gene construction**

| Plasmids | Oligonucleotides | 5'3' Sequence |
|---|---|---|
| pGEM *sec31A1249V::AFpyrG* / pGEM *sec31A1249P::AFpyrG* / pGEM *sec31A1249S::AFpyrG* / pGEM *sec31A1249G::AFpyrG* | sec31 ORF SphI F | GCGGCATGCCGTCCAGTGCACCCATTTCCTC |
| | sec31 3' UTR XbaI R | CGCGTCTAGACGTAATTCCTCGGGAGCGCG |
| | sec31 3' XbaI F | CGCGTCTAGACGGATGAAACGACACTGTCGACG |
| | sec31 3' NotI R | CGCGCGGCCGCCGGCGATCGTTCAACACTGCTAG |
| | AFpyrG XbaI F | CGCGTCTAGAGCCTCAAACAATGCTCTTCACCC |
| | AFpyrG XbaI R | CGGTCTAGACTGTCTGAGAGGAGGCACTGATGCG |
| | sec31 A1249V F | GCACGTGACTACGAGACGGTTCGTACAATCCACATTGA TATCATGAC |
| | sec31 A1249V R | GTCATGATATCAATGTGGATTGTACGAACCGTCTCGTA GTCACGTGC |
| | sec31 A1249P F | GCACGTGACTACGAGACGCCTCGTACAATCCACATTGA TATCATGAC |
| | sec31 A1249P R | GTCATGATATCAATGTGGATTGTACGAGGCGTCTCGTA GTCACGTGC |
| | sec31 A1249S F | GCACGTGACTACGAGACGAGTCGTACAATCCACATTGA TATCATGAC |
| | sec31 A1249S R | GTCATGATATCAATGTGGATTGTACGACTCGTCTCGTAG TCACGTGC |
| | sec31 A1249G F | GCACGTGACTACGAGACGGGTCGTACAATCCACATTGA TATCATGAC |
| | sec31 A1249G R | GTCATGATATCAATGTGGATTGTACGACCCGTCTCGTAG TCACGTGC |
| | sec31 3' check R | CCGAACACAGAGAGCCGAGG |
| | sec31 seq F | GGCCAAGCACAGCCTCTTCTG |
| pGEM *sec16-mCherry* | sec16-ORF-AatII-F | CGCGGACGTCGGTCGTGGATTTCGAAACCAAGCATG |
| | sec16-ORFns-SpeI-R | CGCGACTAGTTTGGGCCATTACATCAACATACCGGC |
| | sec16-3-SpeI-F | CGCGACTAGTGAAGCTCTTAACATCGCTTCCAACTC |
| | sec16-3-NotI-R | CGCGGCGGCCGCGAGGTCCCGCTTCCTGCACTTCAC |
| | mCherry-SpeI-F | GCGACTAGTGGAGCAGGTGCTGGTGCTG |
| | mCherry-XbaI-R-3 | CGCGTCTAGACCTGTTATCCCTAGCGGATCTG |
| pGEM *$thiA_p$-ykt6::AFpyrG* / pGEM *$thiA_p$-ykt6::AFriboB* | ykt6 5' ApaI F | CGCGGGGCCCCTGCAATTGCCTGCATCTGTGCTG |
| | ykt6 5' SpeI R | CGCGACTAGTGCTGGATGAGGCAGGGAGATAATAG |
| | ykt6 ORF SpeI F | CGCGACTAGTATGAAGATCGTTTACATTGGTGTAAGCT GC |
| | ykt6 ORF NotI R | CGCGGCGGCCGCCTACTACTATGGTTGGCGGGATACGG |
| | AFpyrG SpeI F | CGCGACTAGTGCCTCAAACAATGCTCTTCACCCCTC |

| Plasmids | Oligonucleotides | 5'3' Sequence |
|---|---|---|
| | AFriboB SpeI F | CGCGACTAGTCCCGGGCTGCAGGAATTCG |
| | thiAp SpeI R | CGCGACTAGTGTTGACTCAGTTCAATGGTTCGAC |
| pGEM *Δsec22::AFpyrG* | sec22 5' SphI F | CGCGGCATGCGTGCTGTAGGCCCTTGAATGTCC |
| | sec22 5' SpeI R | CGCGACTAGTGTTATAGCTGGGTTCGTTAACTGGCGG |
| | sec22 3' SpeI F | CGCGACTAGT CCCAGCGATGCCACGTAACTACC |
| | sec22 3' NotI R | CGCGGCGGCCGCGGCTCCTTCCAAGTTGTCTCATCG |
| | AFpyrG SpeI F | CGCGACTAGTGCCTCAAACAATGCTCTTCACCCTC |
| | AFpyrG SpeI R | CGGACTAGTCTGTCTGACAGGAGGCACTGATGCG |
| pGEM *thiA$_p$-sec9::AFpyrG* / pGEM *thiA$_p$-FLAG-sec9::AFpyrG* | sec9 5' SphI F | GCGGCATGCCGGAGTTGACTGAACAGTTGGTCAG |
| | sec9 5' SpeI R | CGACTAGTCGACTCCACTAAGCCGAAAGAAGC |
| | sec9 ORF SpeI F | GCGACTAGTATGAAGCGATTCGGCCTCAAAAAGTC |
| | sec9 ORF NotI R | CGCGCGGCCGCGCCAGGCAGTCGTCATAACAATCG |
| | AFpyrG XbaI F | CGCGTCTAGAGCCTCAAACAATGCTCTTCACCC |
| | AFpyrG SpeI R | CGGACTAGTCTGTCTGACAGGAGGCACTGATGCG |
| | thiAp SpeI F | CGCGACTAGTCGACCTGGCACCTACAGAAGAATC |
| | thiAp SpeI R | CGCGACTAGTGTTGACTCAGTTCAATGGTTCGAC |
| | thiApFLAG XbaI R | CGCGTCTAGACTTGTCATCGTCGTCCTTGTAGTCC |
| pGEM *ΔsynA::AFriboB* | synA 5' ApaI F | CGCGGGGCCCGACTTGCCGTGATCATATATCTTGTCC |
| | synA 5' SpeI R | CGCGACTAGTCACGGGTTGTGGAAGAAGAGGAGG |
| | synA 3' SpeI F | CGCGACTAGTCTACGACACTTCAACGACGGTCATG |
| | synA 3' NotI R | CGCGGCGGCCGCGACTTGGTGGTGGAAGCTCTCCG |
| | AFriboB SpeI F | CGCGACTAGTCCCGGGCTGCAGGAATTCG |
| | AFriboB SpeI R | CGCGACTAGTCCCGGGCTGCAGGAATTCGA |
| pGEM *ΔnyvA::AFpyrG* / pGEM *ΔnyvA::AFpyroA* | nyvA 5' ApaI F | CGCGGGGCCCGCTGGGTTCGTTGGATGATGAC |
| | nyvA 5' XbaI R | CGCGTCTAGAGCCAAAATACCAAGAGCCAACCG |
| | nyvA 3' XbaI F | CGCGTCTAGAGGCCATATATGTGCAATTGTGTCAC |
| | nyvA 3' NotI R | CGCGCGGCCGCGTCATGTGACGCACATTCCACTG |
| | AFpyrG XbaI F | CGCGTCTAGAGCCTCAAACAATGCTCTTCACCC |
| | AFpyrG XbaI R | CGGTCTAGACTGTCTGAGAGGAGGCACTGATGCG |
| | AFpyroA SpeI F | CGCGACTAGTGGACATCAGATGCTGGATTAC |
| | AFpyroA SpeI R | CGCGACTAGTGCGAGTGTCTACATAATGAAGG |
| pGEM *alcA$_p$-mCherry-synA::AFpyroA* | AFpyroA XbaI F | CGCGTCTAGAGGACATCAGATGCTGGATTAC |
| | AFpyroA SpeI R | CGCGACTAGTGCGAGTGTCTACATAATGAAGG |
| | alcAp XbaI F | CGCTCTAGATAAGTCCCTTCGTATTTCTCCGC |
| | alcAp XbaI R | CGCGTCTAGAATTTTGAGGCGAGGTGATAGG |
| | mCherry ORF SpeI F | GCGCACTAGTATGGTGAGCAAGGGCGAGG |
| | mCherry NS GA XbaI R | GCGCTCTAGATGCTCCCTTGTACAGCTCGTCCATGCC |
| | 5 AN8769 ApaI F | GCGCGGGCCCCCTGCGAGATCCAATGACTC |
| | 5 AN8769 SpeI R | GCGCACTAGTGATGCTCTGTGCGAAGAGCTTG |
| | AN8769 5 check | CTGATGAATTCAACCGTAGTGTCCG |
| | AN8769 ORF SpeI F | CGCGACTAGTATGTCTGAGCAACCGTACGATCC |
| | AN8769 ORF3 NotI R | CGCGGCGGCCGCGTGACTCAGACCACCAAGATCACC |

| Plasmids | Oligonucleotides | 5'3' Sequence |
|---|---|---|
| pGEM *thiA$_p$-FLAG-sarA::AFpyrG* | AN0411 5 ApaI F | CGCGGGGCCCGAAGACTCGGAAGCAACTAGGTTTC |
| | AN0411 5 SpeI R | CGCGACTAGTCGCCGTGTAAATGCAGACACAAAG |
| | AN0411 ORF SpeI F | CGCGACTAGTATGTGGATTATCAACTGGTGTAAGTCTGC |
| | AN0411 3 NotI R | CGCGCGGCCGCGGAGCTTAACCAGAATGCCAGTG |
| | AFpyrG XbaI F | CGCGTCTAGAGCCTCAAACAATGCTCTTCACCCTC |
| | AFpyrG SpeI R | CGGACTAGTCTGTCTGAGAGGAGGCACTGATGCG |
| | thiAp XbaI F | CGCGTCTAGACGACCTGGCACCTACAGAAGAATC |
| | thiA FLAG XbaI R | CGCGTCTAGACTTGTCATCGTCGTCCTTGTAGTCCATGTTGACTCAGTTCAATGGTTCGACTATAG |
| pGEM *thiA$_p$-FLAG-sec12::AFpyrG* | AN11127 5 ApaI F | CGCGGGGCCCGCGTGTCCGAGAATTTCTGATGG |
| | AN11127 5 XbaI R | CGCGTCTAGACCAGGGGTGTGACAACTAATTGC |
| | AN11127 5 check | GCCAAATACGACATCAGAAACCC |
| | AN11127 ORF XbaI F | CGCGTCTAGAATGGCGCCCAAAATACCGTCTGC |
| | AN11127 ORF NotI R | CGCGGCGGCCGCCAAGATACGGAGGCATGACGCCTC |
| | AFpyrG XbaI F | CGCGTCTAGAGCCTCAAACAATGCTCTTCACCCTC |
| | AFpyrG SpeI R | CGGACTAGTCTGTCTGAGAGGAGGCACTGATGCG |
| | thiAp XbaI F | CGCGTCTAGACGACCTGGCACCTACAGAAGAATC |
| | thiA FLAG XbaI R | CGCGTCTAGACTTGTCATCGTCGTCCTTGTAGTCCATGTTGACTCAGTTCAATGGTTCGACTATAG |
| pGEM *thiA$_p$-FLAG-sec24::AFpyrG* | AN3720 5 SphI F | CGCGGCATGCCGGCTCAAACAAGTCGAAGACCTTATC |
| | AN3720 5 SpeI R | CGCGACTAGTCTAGGCATTTGCAGCTTATGAACGTTTC |
| | AN3720 ORF SpeI F | CGCGACTAGTATGGCATCTCCACAAGGGGGCTAC |
| | AN3720 ORF NotI R | CGCGGCGGCCGCCACAGACACTTGTGCCTTGGAGCAC |
| | AFpyrG XbaI F | CGCGTCTAGAGCCTCAAACAATGCTCTTCACCCTC |
| | AFpyrG SpeI R | CGGACTAGTCTGTCTGAGAGGAGGCACTGATGCG |
| | thiAp XbaI F | CGCGTCTAGACGACCTGGCACCTACAGAAGAATC |
| | thiA FLAG XbaI R | CGCGTCTAGACTTGTCATCGTCGTCCTTGTAGTCCATGTTGACTCAGTTCAATGGTTCGACTATAG |
| pGEM *thiA$_p$-FLAG-sec13::AFpyrG* | AN4317 5 ApaI F | CGCGGGGCCCCCAGCTTGCGTACATCGAATCATG |
| | AN4317 5 SpeI R2 | CGCGACTAGTGAGCTATTTGCAGCTCTAGGCCGATG |
| | AN4317 ORF SpeI F | CGCGACTAGTATGGTACGTCTTCGCCGTCAATTC |
| | AN4317 3 NotI R | CGCGCGGCCGCGGAAGACAGGATTTGCTGGCTTG |
| | AFpyrG XbaI F | CGCGTCTAGAGCCTCAAACAATGCTCTTCACCCTC |
| | AFpyrG SpeI R | CGGACTAGTCTGTCTGAGAGGAGGCACTGATGCG |
| | thiAp XbaI F | CGCGTCTAGACGACCTGGCACCTACAGAAGAATC |
| | thiA FLAG XbaI R | CGCGTCTAGACTTGTCATCGTCGTCCTTGTAGTCCATGTTGACTCAGTTCAATGGTTCGACTATAG |
| pGEM *thiA$_p$-sec31::pabaA*/ pGEM *thiA$_p$-FLAG-sec31::AFpyrG* | 5 sec31 ApaI F | GCGCGGGCCCGTACGTCCGTACAGCTCGCTC |
| | 5 sec31 SpeI R | GCGCACTAGTGCTGAGAGTAGGGGTCTGC |
| | ORF sec31 SpeI F | GCGCACTAGTATGGTGCGTCTGAGGGAGATTC |
| | ORFsec31NotIR | GCGCGCGGCCGCCCTGGGCAAATAATCAACAGTTTC |
| | AFpyrG XbaI F | CGCGTCTAGAGCCTCAAACAATGCTCTTCACCCTC |
| | AFpyrG SpeI R | CGGACTAGTCTGTCTGAGAGGAGGCACTGATGCG |
| | thiAp XbaI F | CGCGTCTAGACGACCTGGCACCTACAGAAGAATC |

| Plasmids | Oligonucleotides | 5'3' Sequence |
|---|---|---|
| | thiA FLAG XbaI R | CGCGTCTAGACTTGTCATCGTCGTCCTTGTAGTCCATGTTGACTCAGTTCAATGGTTCGACTATAG |
| | thiA SpeI F | CGCGACTAGTCGACCTGGCACCTACAGAAGAATC |
| | thiA XbaI R | CGCGTCTAGAGTTGACTCAGTTCAATGGTTCGAC |
| pGEM $thiA_p$*-FLAG-ykt6::AFpyrG* | AN8488 5 ApaI F | CGCGGGGCCCCTGCAATTGCCTGCATCTGTGCTG |
| | AN8488 5 SpeI R | CGCGACTAGTGCTGGATGAGGCAGGGAGATAATAG |
| | AN8488 ORF SpeI F | CGCGACTAGTATGAAGATCGTTTACATTGGTGTAAGCTGC |
| | AN8488 ORF NotI R | CGCGGCGGCCGCCTACTACTATGGTTGGCGGGATACGG |
| | AFpyrG XbaI F | CGCGTCTAGAGCCTCAAACAATGCTCTTCACCCTC |
| | AFpyrG SpeI R | CGGACTAGTCTGTCTGAGAGGAGGCACTGATGCG |
| | thiAp XbaI F | CGCGTCTAGACGACCTGGCACCTACAGAAGAATC |
| | thiA FLAG XbaI R | CGCGTCTAGACTTGTCATCGTCGTCCTTGTAGTCCATGTTGACTCAGTTCAATGGTTCGACTATAG |
| pGEM $thiA_p$*-FLAG-ssoA::AFpyrG* | AN3416 5 ApaI F | CGCGGGGCCCGTGTGTTTTCCCAGGCTCTGGGC |
| | AN3416 5 SpeI R | CGCGACTAGTCAGCTCCTGACGACAGCAGTGAC |
| | AN3416 ORF SpeI F | CGCGACTAGTATGAGTGTATGTCCGCCGGCCTAATG |
| | AN3416 3 NotI R | CGCGGCGGCCGCCGTCAAGCCGCCTCATAGATGCAAC |
| | AFpyrG XbaI F | CGCGTCTAGAGCCTCAAACAATGCTCTTCACCCTC |
| | AFpyrG SpeI R | CGGACTAGTCTGTCTGAGAGGAGGCACTGATGCG |
| | thiAp XbaI F | CGCGTCTAGACGACCTGGCACCTACAGAAGAATC |
| | thiA FLAG XbaI R | CGCGTCTAGACTTGTCATCGTCGTCCTTGTAGTCCATGTTGACTCAGTTCAATGGTTCGACTATAG |
| pGEM *ΔrabD::AFpyrG* | AN6974 5 ApaI F | CGCGGGGCCCGACCTACGAAGTCCTTTTATGGGC |
| | AN6974 5 SpeI R | CTATAATCAGCCGACTAGTGGTTCG |
| | AN6974 3 NotI R | CGCGGCGGCCGCCTCCGGCGTTTCTACCGATTG |
| | AN6974 3 SpeI F | GGGACTAGTATGTGTATTCTCCTG |
| | AN6974 5 check | GAAACTTGTCCGGACCATACGCC |
| | AFpyrG SpeI F | CGCGACTAGTGCCTCAAACAATGCTCTTCACCCTC |
| | AFpyrG SpeI R | CGGACTAGTCTGTCTGACAGGAGGCACTGATGCG |
| pGEM $thiA_p$*-rabO::AFpyrG* | rabO 5 ApaI F | CGCGGGGCCCGAAGGGTTCAAGGATGAAACGAAC |
| | rabO 5 XbaI R | CGCGTCTAGACTCAGTGCAGGAGAGCAAAGGAG |
| | rabO ORF XbaI F | CGCGTCTAGAATGAACCCTGAGTGGTAAGTGCTTTG |
| | rabO 3 NotI R | CGCGGCGGCCGCCAACCTCTAATTTACCACGGCAGC |
| | AFpyrG SpeI F | CGCGACTAGTGCCTCAAACAATGCTCTTCACCCTC |
| | AFpyrG XbaI R | CGGTCTAGACTGTCTGAGAGGAGGCACTGATGCG |
| | thiAp SpeI F | CGCGACTAGTCGACCTGGCACCTACAGAAGAATC |
| | thiAp SpeI R | CGCGACTAGTGTTGACTCAGTTCAATGGTTCGAC |
